# A human photoacoustic imaging reporter gene using the clinical dye indocyanine green

**DOI:** 10.1101/537100

**Authors:** Nivin N. Nyström, Lawrence C.M. Yip, Jeffrey J.L. Carson, Timothy J. Scholl, John A. Ronald

**Author notes:** **Corresponding Author’s Contact Information:** John Ronald, 1151 Richmond St. N. Room 2241A, Robarts Research Institute, University of Western Ontario, London, ON Canada N6A 5B7.

## Abstract

Photoacoustic imaging (PAI) combines optical contrast with the resolution and depth-detection of ultrasound and is increasingly being utilized for medical imaging in patients. PAI reporter genes would allow for monitoring of cell and gene therapies, but current reporters have immunogenicity and/or toxicity concerns that may limit clinical translation. Here we report a PAI reporter system employing the ability of human *organic anion transporting polypeptide 1b3* (*Oatp1b3*) to take up the clinical dye indocyanine green (ICG) into cells. Following ICG administration, cells synthetically expressing *Oatp1b3* exhibited significantly increased PAI signals compared to control cells both *in vitro* and in mice. Several benefits of this technology are the human derivation of *Oatp1b3*, and the high extinction coefficient, low quantum yield and pre-existing clinical approval of ICG. We posit that the *Oatp1b3*-ICG reporter system could be useful for *in vivo* gene and cell tracking in preclinical and clinical applications.

---

Reporter genes are important tools for detecting cellular and molecular processes *in vivo*^1^, and have contributed extensively to monitoring fates of various cell types such as cancer, immune and stem cells in preclinical models^2, 3, 4, 5, 6^. Clinical studies have also applied reporter gene technology for monitoring of gene^7, 8^ and cell-based^9^ therapies in cancer patients via positron emission tomography (PET) reporters. PET produces whole-body images with high sensitivity, but has limitations such as cost, availability, and ionizing radiation concerns for longitudinal studies. Reporter genes for affordable, portable, and safer imaging modalities are sought for continued monitoring of localized gene and cell tracking in humans. Fluorescence reporters have been united with several *in vivo* preclinical and clinical optical modalities such as intravital microscopy, whole-body small animal fluorescence imaging (FLI), and fluorescence endoscopy^10, 11, 12, 13, 14^. Yet, light attenuation and scattering by surrounding tissues can obscure detection of deeply-located and/or weak fluorescence signals, limiting it to superficial depths in humans^15,16^. Ultrasound (US) acoustic waves, on the other hand, are less susceptible to scattering and attenuation, permitting signal detection at increased depth and resolution^1^.

Photoacoustic imaging (PAI), also called optoacoustic imaging, detects acoustic waves from optically-excited sources, combining benefits of US resolution and depth with optical contrast. Clinical PAI is being applied to a growing number of biomedical applications, spanning from sentinel lymph node imaging to vertebrae imaging for spinal surgery guidance^17, 18, 19, 20, 21, 22, 23, 24^. PAI reporter genes that operate in the near-infrared (NIR) window (650 to 1350 nm) are particularly desirable due to the absence of endogenous chromophores absorbing at these wavelengths^25^. Established PAI reporters include NIR fluorescent proteins^26, 27^; bacterial *LacZ*, which enzymatically generates a blue pigment from the exogenous probe X-gal^28^; and human *tyrosinase*, which initiates the conversion of endogenous tyrosine into melanin^29, 30, 31, 32, 33, 34, 35^. Though established PAI reporters have been useful in preclinical studies, their clinical utility may be limited due to their non-human origin, suboptimal sensitivity and specificity, and/or undesirable toxicity.

Here we report a novel PAI reporter gene called human *organic anion-transporting polypeptide 1b3* (*Oatp1b3*), that operates by taking up the clinically-utilized NIR fluorescent dye indocyanine green (ICG). *Oatp1b3* is endogenously expressed in the human liver and is partly responsible for ICG uptake during liver function tests^36^. Previous work has shown that *Oapt1b3* and ICG can be utilized as a reporter gene-probe pair for FLI^37, 38^. We extend this work by establishing this pair as a PAI reporter system. With ICG as the source of contrast, this system has several benefits for PAI including its human origin, low probe toxicity with longstanding FDA-approval^39^, a high molar extinction coefficient^40^, relatively low fluorescence quantum yield^41^, signal amplification (*i.e*., one transporter takes up multiple ICG molecules), and a narrow, distinct NIR absorption band peaking at 805 nm *in vivo*. This system would be particularly useful for studying whole-body *in vivo* gene expression in preclinical models, and for tracking of gene- and cell-based therapies at both the pre-clinical and clinical stages.

## RESULTS

### Generation of breast cancer cells expressing fluorescent reporters and *Oatp1b3*

Human (MDA-MB-231) and murine (4T1) triple negative breast cancer cell lines were transduced first with lentivirus encoding the fluorescence reporter *tdTomato* and were subsequently sorted for tdTomato fluorescence with >95% purity to obtain *tdTomato*-expressing cells, referred to hereon in as tdTomato Control cells. Following expansion, an aliquot of these cells was additionally transduced with a second lentivirus co-encoding the fluorescence reporter *zsGreen1* (zsG) and *Oatp1b3* (**Figure 1A**). This second cell population was sorted for both tdTomato intensity equivalent to the tdTomato Control cell population, as well as zsGreen intensity with >90% purity to generate tdTomato OATP1B3 cells (**Figure 1B**). Immunofluorescence staining validated absence of *Oatp1b3* expression in tdTomato Control cells, whereas positive staining was present in the tdTomato OATP1B3 cells (**Figure 1C; Supplementary Figure 1A**). Proliferation assays showed no significant difference in growth rates between non-transduced cells, tdTomato Control cells, and tdTomato OATP1B3 cells, or between cell populations incubated in 35-μg/ml ICG for one hour (**Figure 1D; Supplementary Figure 1B**).

**Figure 1.**
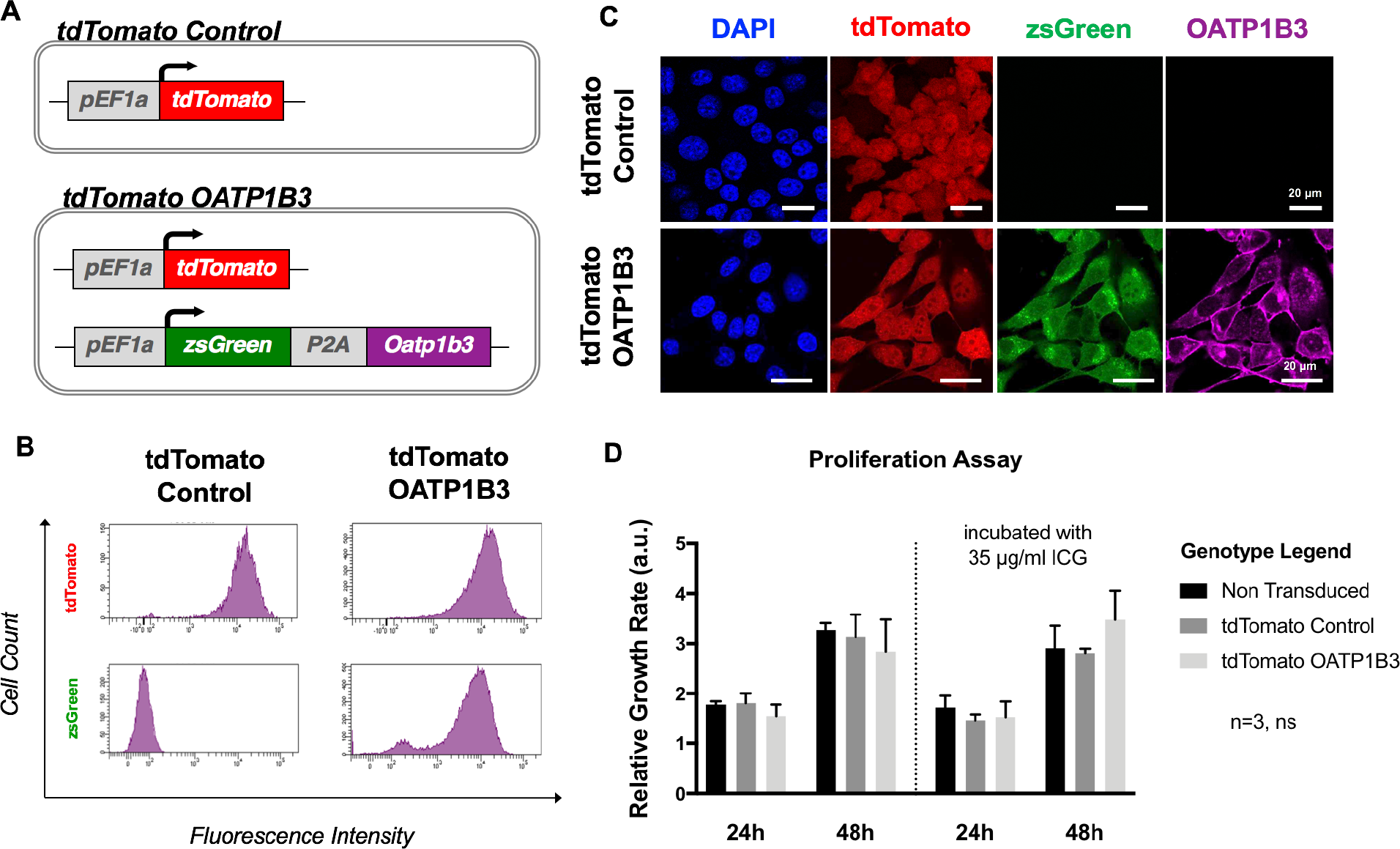
Cell Engineering and Characterization. A) Reporter gene constructs used to engineer cells, which include the fluorescence reporter *tdTomato* (top; tdTomato Control), or *tdTomato* in addition to the fluorescence reporter *zsGreen* (zsG) with *Organic anion-transporting polypeptide 1b3* (*Oatp1b3*) separated by the self-cleaving peptide (P2A) (bottom; tdTomato OATP1B3), each under control of the human elongation factor 1 promoter (*pEF1a*). B) Fluorescence-activated cell sorting plots following lentivirus transduction of MDA-MB-231 cells with tdTomato Control transfer plasmids or tdTomato OATP1B3 transfer plasmids. C) Microscopy of engineered MDA-MB-231 cells for nuclear staining (DAPI; blue), tdTomato fluorescence (red), zsGreen fluorescence (green), and immunofluorescence staining for OATP1B3 expression (purple). Scale bar, 20 μ;m. D) Relative growth rates (arbitrary units, a.u.) of Non Transduced, tdTomato Control, and tdTomato OATP1B3 cells grown in the presence or absence of 35 μ;g/ml indocyanine green (ICG). Error bars represent one standard deviation.

### *In vitro* ICG-enhanced FLI and PAI of breast cancer cells expressing *Oatp1b3*

Next, we evaluated FLI and PAI contrast of both human and murine breast cancer cells expressing *Oatp1b3* incubated with or without ICG by imaging cell pellets in a phantom. Both MDA-MB-231 and 4T1 tdTomato OATP1B3 cells incubated with 35-μg/ml ICG (and subsequently washed) exhibited significantly increased fluorescence radiance (p/s/cm^2^/sr) at ICG wavelengths relative to ICG-incubated tdTomato Control cells (4.0-fold, p<0.05; **Figure 2A**). Fluorescence radiance from ICG-incubated tdTomato OATP1B3 cells was also significantly increased relative to an equivalent volume of 35-μg/ml ICG alone, referred to as the positive control (2.6-fold; p<0.05). No significant difference in fluorescence radiance at ICG wavelengths was observed between tdTomato Control and tdTomato OATP1B3 cells not incubated with ICG, ruling out any contribution of zsGreen or OATP1B3 to signals at NIR wavelengths. PAI of the same phantom at 780 nm was acquired using a custom-built PAI system. Photoacoustic contrast-to-noise ratio (CNR; arbitrary units, a.u.) was significantly increased for both MDA-MB-231 (2.7-fold; p<0.05) and 4T1 (2.4-fold; p<0.05) tdTomato OATP1B3 cells incubated with 35 μg/ml-ICG, relative to CNR of ICG-incubated tdTomato Control cells and all untreated controls (**Figure 2B**). PAI of tdTomato Control cells incubated with ICG did not exhibit a significant difference in CNR relative to untreated control cells. Further, *Oatp1b3* has been previously established as a magnetic resonance imaging (MRI) reporter gene, based on its ability to take up the positive contrast, clinically-utilized paramagnetic contrast agent gadolinium ethoxybenzyl diethylenetriaminepentaacetic acid (Gd-EOB-DTPA)^37^. To evaluate this in our cells, we performed inversion recovery magnetic resonance imaging of tdTomato Control and tdTomato OATP1B3 MDA-MB-231 cells on a 3-Tesla MRI scanner and found significantly increased spin-lattice relaxation rates (4.9-fold, p<0.05) from tdTomato OATP1B3 cells incubated with 6.4 mM Gd-EOB-DTPA, compared to all other control conditions (**Supplementary Figure 2**).

**Figure 2.**
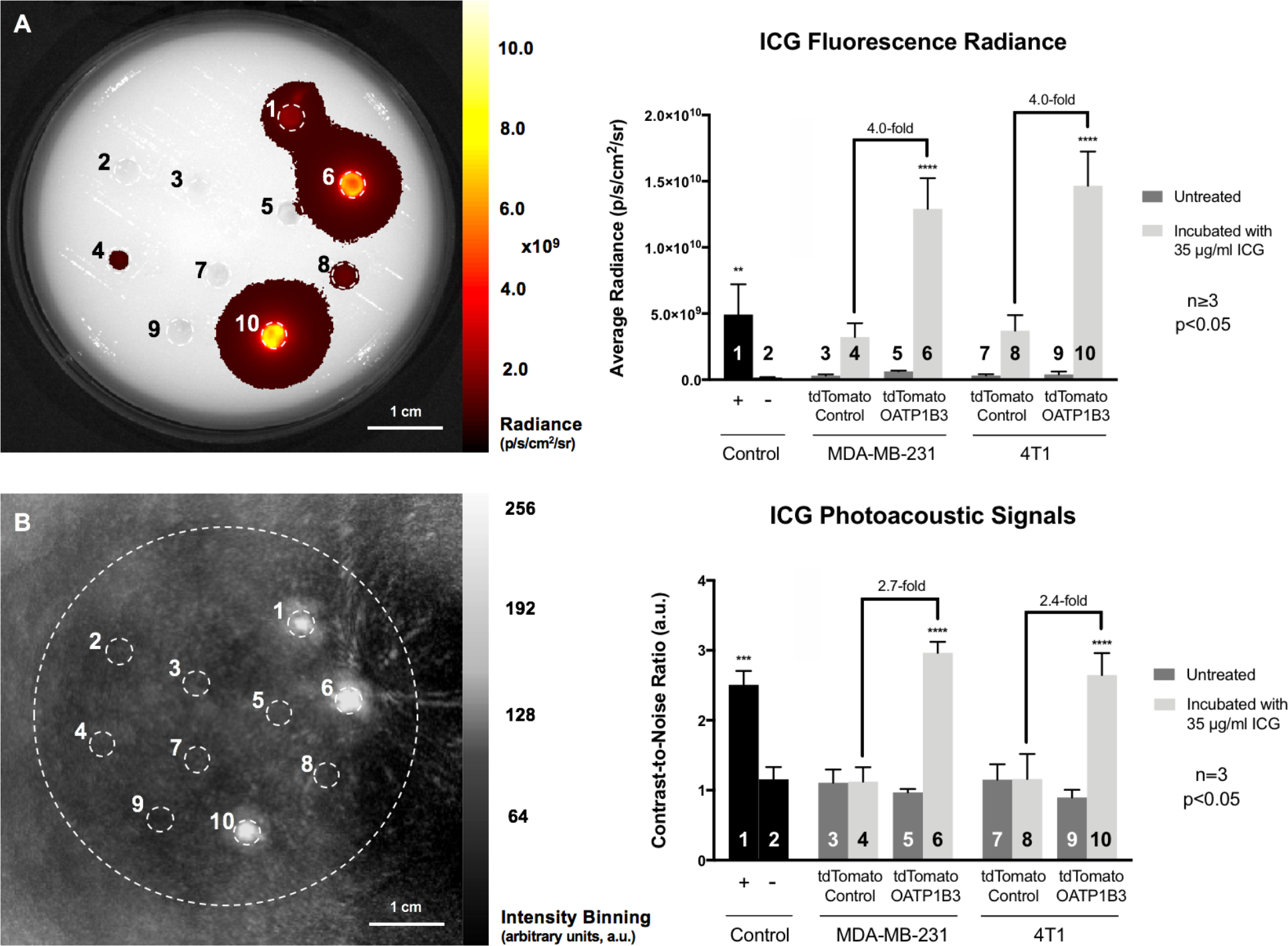
*In Vitro* Fluorescence and Photoacoustic Imaging. A) Fluorescence imaging (FLI) acquired with 780-nm excitation and 845-nm emission filters of an asymmetrical custom-built phantom with well containing: (*1*) 35-μg/ml indocyanine green (ICG) in media; (*2*) media alone; (*3*) human breast cancer (MDA-MB-231) tdTomato Control cells; (*4*) MDA-MB-231 tdTomato Control cells incubated with 35 μg/ml ICG; (*5*) MDA-MB-231 tdTomato OATP1B3 cells; (*6*) MDA-MB-231 tdTomato OATP1B3 cells incubated with 35-μg/ml ICG; (*7*) murine breast cancer (4T1) tdTomato Control cells; (*8*) 4T1 tdTomato Control cells incubated with 35-μg/ml ICG; (*9*) 4T1 tdTomato OATP1B3 cells; and (*10*) 4T1 tdTomato OATP1B3 cells incubated with 35-μg/ml ICG. Fluorescence radiance (p/s/cm^2^/sr) is quantified for each well region of interest. B) Photoacoustic imaging (PAI) of the same phantom acquired with an optical excitation at 780 nm. Contrast-to-noise ratios (relative to the center of the phantom) were quantified (arbitrary units, a.u.). Numbers, *1-10*, on graphs correspond to numbers on images. Scale bar, 1 cm. Error bars represent one standard deviation.

### ICG-enhanced FLI signal from *Oatp1b3*-expressing tumors

The fourth mammary fat pad of immunocompromised female mice was implanted with tdTomato Control (n=4) or tdTomato OATP1B3 (n=4) MDA-MB-231 cells to generate orthotopic tumors for *in vivo* FLI and PAI. FLI for tdTomato activity was performed (5-min exposure time; 520-nm excitation filter, 570-nm emission filter) to obtain relative measures of the total number of viable cells for each tumor. No significant difference in tdTomato radiance (p/s/cm^2^/sr) was found between tdTomato Control and tdTomato OATP1B3 tumors over time (**Supplementary Figure 3A**). Longitudinal FLI for ICG fluorescence (5-min exposure time; 780-nm excitation filter, 845-nm emission filter) after administration of 8-mg/kg ICG determined that the greatest significant difference in ICG signals between tdTomato Control and tdTomato OATP1B3 tumors occurred at 24 hours post-ICG (**Supplementary Figure 3B**). Accordingly, a second cohort of animals was imaged prior to ICG administration and 24 hours post-ICG. Neither tdTomato Control (n=6) nor tdTomato OATP1B3 (n=5) tumors generated FLI radiance at ICG wavelengths prior to ICG administration, confirming no possible contribution by zsGreen or OATP1B3 proteins to signals detected at ICG wavelengths (**Figure 3A**, *middle column*). Prior to ICG administration, fluorescence measures for ICG were not significantly different between tdTomato Control tumors (9.14 ± 3.49 × 10^6^ p/s/cm^2^/sr) and tdTomato OATP1B3 tumors (7.41 ± 1.86 × 10^6^ p/s/cm^2^/sr). Twenty-four hours after ICG administration (8 mg/kg), tdTomato OATP1B3 tumors exhibited significantly increased (8.9-fold) fluorescence (7.36 ± 1.77 × 10^8^ p/s/cm^2^/sr) relative to tdTomato Control tumors (8.22 ± 0.527 × 10^7^ p/s/cm^2^/sr; p<0.05; **Figure 3B**). FLI signals at ICG wavelengths were also detected from the abdomen of both tdTomato Control and tdTomato OATP1B3 tumors 24 hours following ICG administration, likely from physiological hepatobiliary clearance of ICG via the gastrointenstinal system^42^ (**Figure 3A**, *last column*).

**Figure 3.**
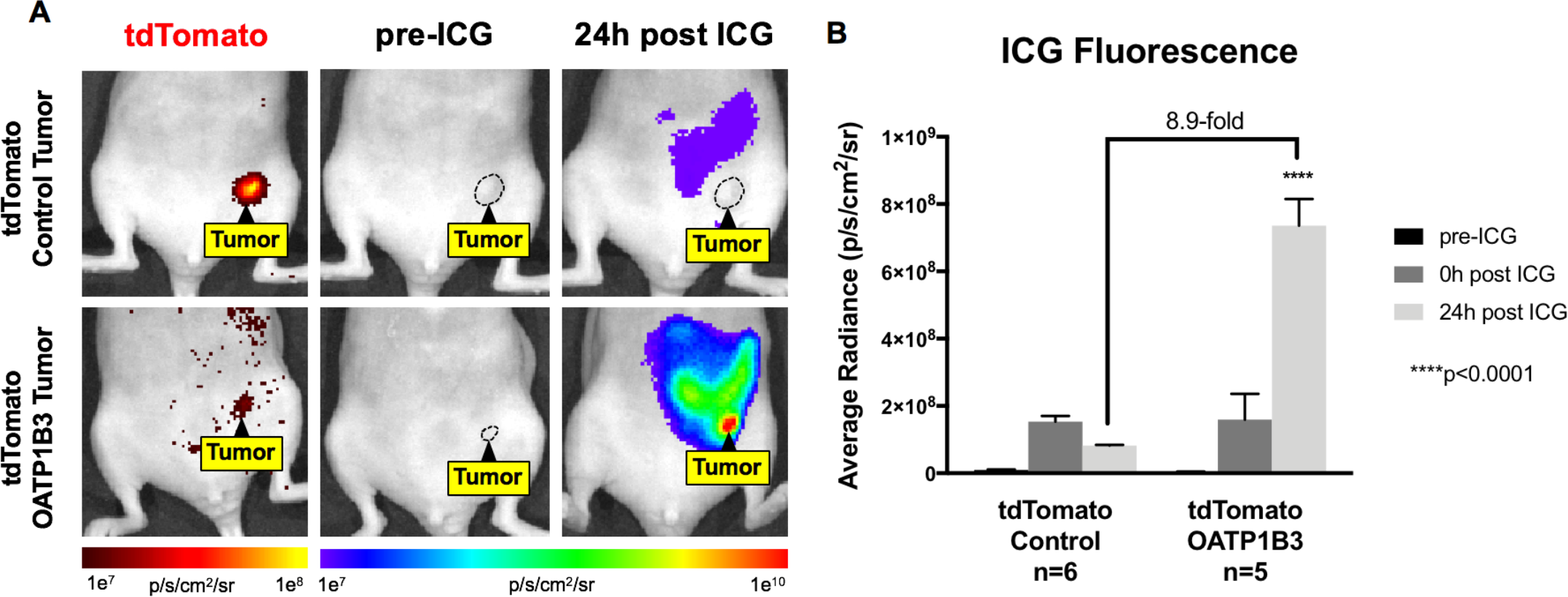
*In Vivo* Fluorescence Imaging of Engineered Tumors. A) Fluorescence imaging (FLI) for tdTomato (*first column*) and indocyanine green (ICG) average radiance (p/s/cm^2^/sr) before (pre-ICG; *second column*) and 24 hours following administration of 8-mg/kg ICG (24h post ICG; *third column*) of a representative mouse burdened with a human breast cancer (MDA-MB-231) tdTomato Control tumor (*top panel*) or with a MDA-MB-231 tdTomato OATP1B3 tumor (*bottom panel*). Tumors are outlined with black dashed lines and/or pointed out via the “Tumor” label. tdTomato signal (*first column*) is directly proportional to tumor size. B) Fluorescence radiance (p/s/cm^2^/sr) of tdTomato Control and tdTomato OATP1B3 tumors before administration of indocyanine green (ICG; pre-ICG), immediately following intraperitoneal injection of ICG (0h post ICG), and 24 hours following ICG administration (24h post ICG). Error bars represent one standard deviation.

### ICG-enhanced PAI signal from *Oatp1b3*-expressing tumors

To evaluate OATP1B3-ICG as a PAI reporter system, mice were also imaged pre- and post-contrast on a small animal PAI scanner (Vevo LAZR-X system Fujifilm Visual Sonics). Both US and NIR-spectrum (680-970nm; Δλ=5nm) PAI of tumors was performed before and 24 hours following intraperitoneal administration of ICG (8 mg/kg). Prior to ICG administration, tdTomato Control and tdTomato OATP1B3 tumor morphology was discernible via US, enabling acquisition of PAI baseline signals at field of views encompassing tumor boundaries (**Figure 4A**; i, iii). Twenty-four hours following ICG administration, tdTomato OATP1B3 tumors demonstrated qualitative increases in signal intensity relative to pre-contrast images; tdTomato Control tumors did not exhibit this increase (**Figure 4A**; ii, iv). Importantly, tdTomato OATP1B3 tumors exhibited increased PAI signal within a narrow absorption band spanning from 695 to 855 nm, with a consensus maximum peak at 805 nm across all tdTomato OATP1B3 tumors 24 hours post-ICG. By comparison, tdTomato Control tumors did not exhibit any increases in PA signal within any region of the NIR spectrum following ICG injection, such that NIR spectra prior-to and 24 hours after ICG administration appeared equivalent (**Figure 4B**). Since there was variance in tumor size, background signals, and tumor morphology within and across tdTomato Control and tdTomato OATP1B3 tumor groups, mean hemoglobin (HbO_2_) PA signals, measured via acoustic average (a.u.) as λ ∈ (855, 955); Δλ=5nm, was calculated for each acquired spectrum and used to normalize the signal acquired at 805 nm. This ICG-to-HbO_2_ signal ratio, referred to as normalized PA signal (a.u.), was not statistically different between tdTomato Control (0.72 ± 0.09 a.u.) and tdTomato OATP1B3 (0.85 ± 0.16 a.u.) tumors prior to ICG injection. Normalized PA signal was also not significantly different between images of tdTomato Control tumors collected before and after ICG administration (0.75 ± 0.13 a.u.; **Figure 4C**). In contrast, 24 hours post ICG, tdTomato OATP1B3 tumors demonstrated significantly increased normalized PA signals (1.92 ± 0.40 a.u.) relative to all other groups (p<0.05).

**Figure 4.**
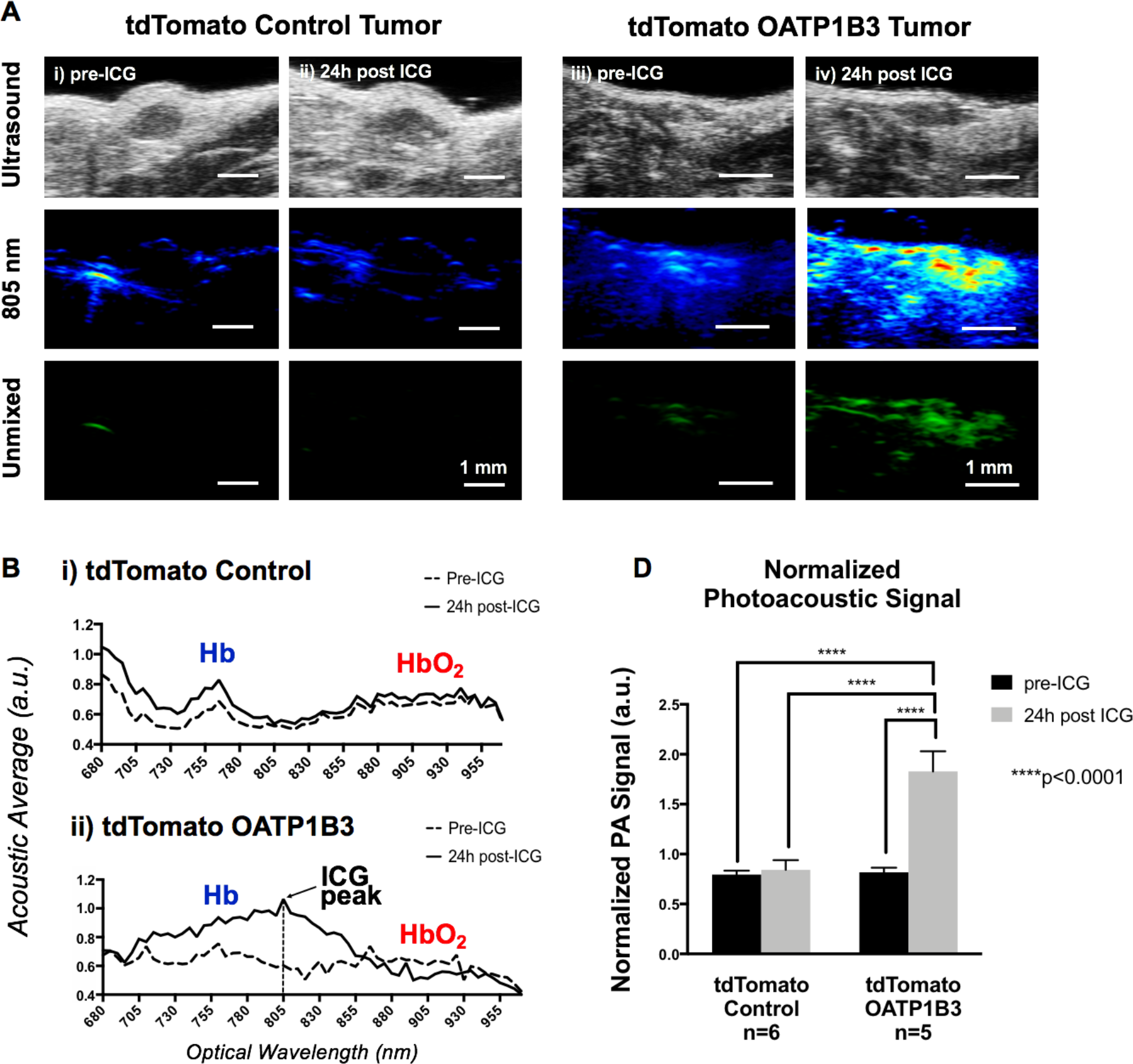
*In Vivo* Near-Infrared Spectral Photoacoustic Imaging. A) Ultrasound image (*top row*), PAI image at 805 nm (*second row*), and a spectrally unmixed image of indocyanine green (ICG) distribution across all wavelengths (Unmixed; *third row*) of a representative human breast cancer (MDA-MB-231) tdTomato Control Tumor (*left panel*) and tdTomato OATP1B3 Tumor (*right panel*), both prior-to administration of ICG (i, iii; pre-ICG) and 24 hours following administration of 8-mg/kg ICG (ii, iv; 24h post ICG). PAI contrast enhancement on PAI is exhibited by tdTomato OATP1B3 tumors 24 hours following ICG administration, while tdTomato Control tumors do not exhibit increased signals 24 hours following ICG administration. B) Near-infrared PAI spectra of a representative tdTomato Control Tumor (*top*) and tdTomato OATP1B3 tumor (*bottom*), before administration of ICG (*dashed-line plot*) and 24 hours following administration of 8-mg/kg ICG (*solid-line plot*), each labelled with hemoglobin (HbO_2_; red) and deoxyhemoblobin (Hb; blue) peaks. Increased photoacoustic signals are observed exclusively within tdTomato OATP1B3 tumors 24h post ICG, with a maximum at 805 nm (ICG peak). C) Normalized PA signal (Signal*805*/Signal*hemoglobin*) across tumor and treatment groups. Scale bar, 1 mm. Error bars represent one standard deviation.

### *Ex vivo* FLI confirms selective ICG retention in *Oatp1b3*-expressing tumors

Following *in vivo* PAI, mice were sacrificed and selective retention of ICG in tdTomato OATP1B3 tumors, relative to tdTomato Control tumors, was validated via *ex vivo* FLI of 150-μ;m tumor sections (**Figure 5A**). tdTomato fluorescence (5-s exposure, 520-nm excitation filter, 570-nm emission filter) was successfully detected from both tdTomato Control and tdTomato OATP1B3 tumor sections (**Figure 5A**, *first column*). As expected, zsG fluorescence (10-s exposure, 480-nm excitation filter, 520-nm emission filter) was absent in tdTomato Control tumor sections but present in tdTomato OATP1B3 tumor sections (**Figure 5A**, *middle column*). Importantly, ICG FLI signal (10-s exposure, 780-nm excitation filter, 845-nm emission filter) was visible only in tdTomato OATP1B3 tumor sections (**Figure 5**, *final column*). Fluorescence microscopy was also conducted, and further validated the presence and/or absence of tdTomato and zsGreen signals from FLI of tumors (**Figure 5B**).

**Figure 5.**
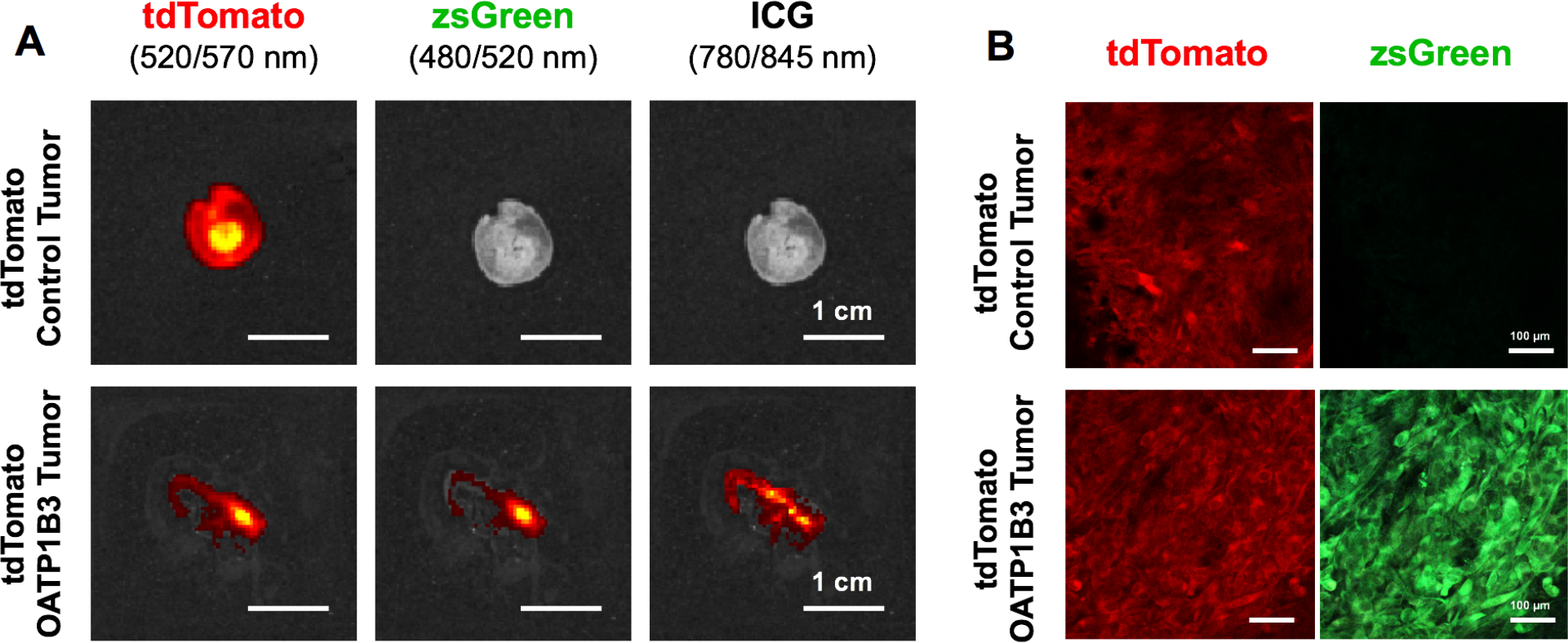
*Ex Vivo* Fluorescence Imaging. A) Fluorescence imaging (FLI) for tdTomato fluorescence (*first column*), zsGreen fluorescence (*second column*), and indocyanine green (ICG; *third column*) of tdTomato Control (*top row*) and tdTomato OATP1B3 (*second row*) tumor slices. Scale bar, 1 cm. B) Fluorescence microscopy for tdTomato (*first column*) and zsGreen (*second column*) of tdTomato Control (*top row*) and tdTomato OATP1B3 (*second row*) tumor slices. Scale bar, 100 μ;m.

## DISCUSSION

In this study we establish *Oatp1b3* and ICG as a novel reporter gene-probe system for *in vivo* PAI of engineered cell populations. We generated a novel lentiviral vector encoding *Oatp1b3*, allowing for engineering of cell lines to stably express this ICG transporter. Cells engineered to express *Oatp1b3* demonstrated significantly increased fluorescence signals at NIR wavelengths following incubation with ICG, which was not observed in control cells. On PAI, *Oatp1b3*-expressing cells incubated with ICG exhibited significantly increased signals *in vitro*, while ICG-incubated control cells were virtually undetectable. We also demonstrated that *Oatp1b3*-expressing breast cancer cells can be implanted into animals to grow orthotopic tumors, and significantly increased FLI and PAI signal are exhibited by small tumors (<1mm^3^) 24 hours following administration of ICG. Importantly, we demonstrate that the increase in PAI detection was specific to the 695 to 855-nm wavelength range, with a consensus maximal peak at 805 nm, allowing for accurate *in vivo* identification of reporter gene-enhanced PAI signal. We further validate the selective retention of ICG exclusively by *Oatp1b3*-expressing tumors using *ex vivo* FLI of tumor slices.

Recently, *Oatp1b3* had been shown to act as a dual MRI and FLI reporter gene system, due to its ability to uptake the clinically-used paramagnetic agent Gd-EOB-DTPA and ICG, respectively^37^. Our study extends this work, establishing *Oatp1b3* as a multimodality reporter for numerous clinically-relevant modalities such as FLI (**Figure 3**), and now PAI (**Figure 4**), and also introduces potential for *Oatp1b3* reporter gene imaging on a clinical 3-Tesla MR scanner (**Supplementary Figure 3**). Extending this system to PAI is important as PAI uniquely offers a non-ionizing and cost-effective method to detect deeply-located engineered cell populations with sensitivity, specificity, and high resolution at sites that are not hindered by optically-scattering materials, though more penetrative PA systems are continuously being developed for such sites^43, 44, 45, 46^. It is important to caution, however, that ICG is endogenously taken up and cleared by the liver and kidneys. Detection of engineered cell populations at these sites, where clearance pathways may limit retention of ICG by *Oatp1b3*-expressing cells, could be challenging. Notwithstanding, the ability to evaluate the status of a gene or cell population using whole-body (MRI), localized (PAI) and microscopic (FLI) imaging techniques with a single reporter gene system could be broadly useful for many biomedical applications, such as tracking the viability and migration of therapeutic cells in patients^15, 47, 48, 49^. Further to the multimodality potential of this reporter, rat-derived *Oatp1a1* and the tracer ^111^Indium-EOB-DTPA have also been established as a reporter gene-probe pair for single photon emission computed tomography (SPECT)^50^. Studies evaluating *Oatp1b3* as a SPECT (or PET) reporter gene are warranted to further extend the potential utility of this human reporter beyond FLI, PAI, and MRI, and improve detection via the unparalleled sensitivity offered by SPECT/PET.

The ideal PAI reporter gene would be specific to the biological process of interest; exhibit an absorption maximum within the NIR window for deep tissue *in vivo* imaging; and, be nontoxic to the cell. In 2007, the bacterial-derived *LacZ* gene was first described as a PAI reporter gene. It encodes the β-galactosidase enzyme that cleaves an exogenously-administered colourless lactose analog, X-gal, to produce the blue pigment, 5,5′-dibromo-4,4′-dichloroindigo. This system featured an elevated absorption coefficient between 600-700 nm^28^. However, *in vivo* applications of *LacZ*-enhanced photoacoustic imaging remain limited due to high endogenous activity of mammalian β-galactosidase (*GLB1*) in most tissues^51, 52^, in conjunction with difficulties in systemic delivery of X-gal^25^. Further, the non-human origin of this gene increases the likelihood of immunogenic reactions in humans, further impeding its application for clinical utility. By comparison, *Oatp1b3* is directly obtained from the human genome, mitigating immunogenic concerns for use in human patients. Additionally, ICG circulates rapidly and readily through the body enabling relatively easy systemic delivery. ICG is not retained within endogenous tissues, even in cases of pathology, for periods of 24 hours following administration, thus eliminating confounding background signals that may ambiguate data, which in turn optimizes contrast-to-noise and specifies signals to the biological process of interest.

Though other reporter genes for PAI have been described (*e.g.*, NIR fluorescent proteins^53, 54^, photoswitchable fluorescent proteins^55^) clinically-relevant PAI reporter gene development has largely focused on *tyrosinase*, which initiates the conversion of endogenous tyrosine into detectable melanin^29, 30, 31, 32, 33, 34, 35^. Though *tyrosinase* presents several desirable properties, including human origin, signal amplification due to enzymatic action, and a detectible product with high photostability, it is not without limitations^56^. The melanin photoacoustic profile displays a broadband, featureless absorption spectrum, making it difficult to distinguish ectopic melanin from other intrinsic signals with high accuracy^25^. Our OATP1B3-ICG system, on the other hand, features a distinct band within the NIR region that is easy to identify and un-mix from hemoglobin and deoxyhemoglobin spectra, and from spectra of other PA reporters added to the biological system for more complex functional imaging. In addition, altered cell phenotype and toxicity associated with ectopic *tyrosinase* activity has been greatly documented^29, 57, 58^, in so much that *tyrosinase*-encoding plasmid distributors (*e.g.*, Imanis Life Sciences), recommend inducible promoters to minimize undesired biological effects during imaging studies. Though studies are limited in number, ectopic *Oatp1b3* overexpression has not been reported to cause cellular toxicity or confer observable phenotypic changes^38, 59^, and we did not observe any quantitative or qualitative differences in cell proliferation rates and/or morphology in our own work.

Moreover, ICG has been used for fluorescence-enhanced medical procedures since 1959, featuring a rare combination of desirable properties, thus making it difficult to displace in the clinic. First, it has low toxicity^39^. It also has a very short half-life in circulation (~5 min), and accordingly, it is not uncommon to inject multiple consecutive doses during fluorescence-enhanced procedures^39^. For our PAI reporter system, short half-life acts as a further advantage, as non-specific ICG not taken up and retained specifically by *Oatp1b3*-expressing cells has largely cleared from the biological system when we image at 24 hours. Further, pre and post-ICG images can be acquired to compare endogenous background signals prior to contrast-enhancement of *Oatp1b3*-engineered cells. ICG also exhibits a relatively high molar extinction coefficient (2.621×10^5^ M^−1^cm^−1^ at 780 nm) at micromolar concentrations^40^ and a relatively low fluorescence quantum yield (0.027 in H_2_O) that decreases with increasing ICG concentration^41^. Signal amplification is also accomplished, given that one OATP1B3 transporter can influx multiple ICG molecules. In summary, the *Oatp1b3*-ICG reporter system marks an important step forward in clinically-relevant reporter gene-enhanced PAI.

In the clinic, PET reporter genes have so far been used in patients for imaging of adenoviral gene therapy^7, 8^ and cytotoxic T cell immunotherapies^9^ in cancer patients. These studies mark major milestones in reporter gene imaging, but limited access and costs due to associated infrastructure and operation, in combination with concerns over ionizing radiation largely hinder broad applicability and longitudinal imaging via PET reporter genes. These limitations highlight the need for alternative reporter gene systems for specific, safe, cost-effective, and accessible imaging modalities within the clinical environment. PAI is a highly scalable imaging modality – different resolutions and imaging depths are achievable based on configurations of light sources, ultrasonic detection systems, and scanning mechanisms across different pre-clinical and clinical PAI systems^60^. Our described reporter system generates a distinct NIR peak via PAI specifically by engineered cells, high CNR relative to surrounding signals, and low toxicity to the biological system, while retaining benefits such as relative affordability, rapid image acquisition, portability, and safety. Future work focuses on further characterizing this reporter gene system for clinically-relevant modalities such as NIR spectroscopy (NIRS), fluorescence endoscopy, and photoacoustic flow cytometry. We expect that this multimodality reporter system will be highly valuable for tracking of gene- and cell-based therapies at both preclinical and clinical stages.

## ACKNOWLEDGEMENTS

Financial support for this manuscript was provided by Natural Sciences and Engineering Research Council (NSERC) Discovery Grants (JAR RGPIN-2016-05420, JJLC RGPIN-2014-04769, TJS RGPIN-2017-06338), the Canada Foundation for Innovation (CFI) Leaders Opportunity Fund (JJLC 29864), the Canadian Institutes of Health Research (CIHR) Collaborative Health Research Projects (CHRP) (NSERC partnered) (JJLC 356794), a Canadian Institutes of Health Research Project Grant (JAR 377071) and an Ontario Institute for Cancer Research Investigator Award (TJS IA-028). This research was additionally supported by the Breast Cancer Society of Canada (LCMY, NNN). The authors would also like to thank the FujiFilm VisualSonics, Inc. team for allowing us to use their PAI scanner and image analysis software to carry out our animal studies.

---

**Supplementary Figure 1.**
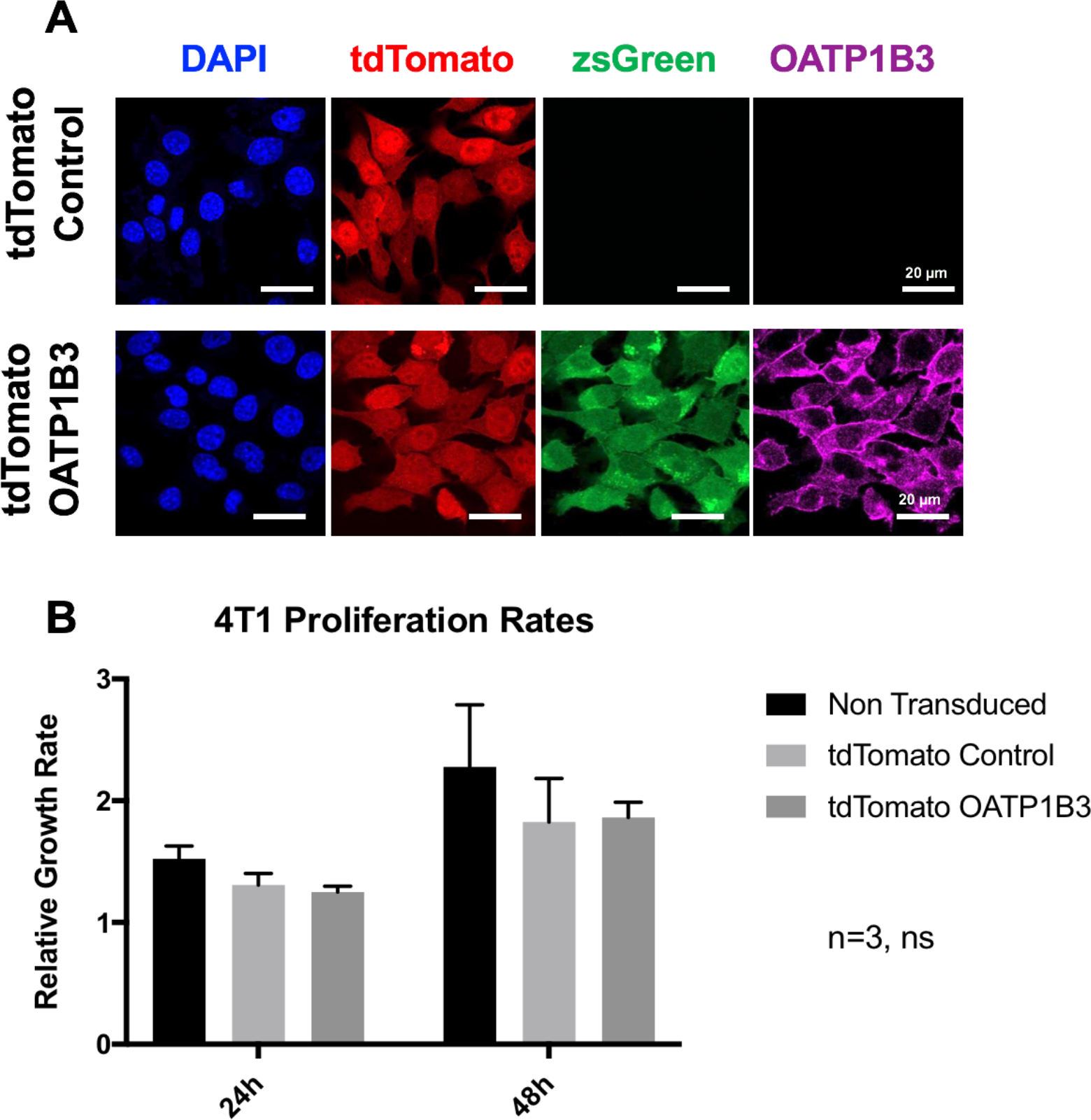
Characterization of Engineered Murine Breast Cancer (4T1) Cells. A) Microscopy of engineered 4T1 cells for nuclear staining (DAPI; blue), tdTomato fluorescence (red), zsGreen fluorescence (green), and immunofluorescence staining for OATP1B3 expression (purple). Scale bar, 20 μ;m. B) Relative growth rates of Non Transduced, tdTomato Control and tdTomato OATP1B3 cells. Error bars show standard deviation.

**Supplementary Figure 2.**
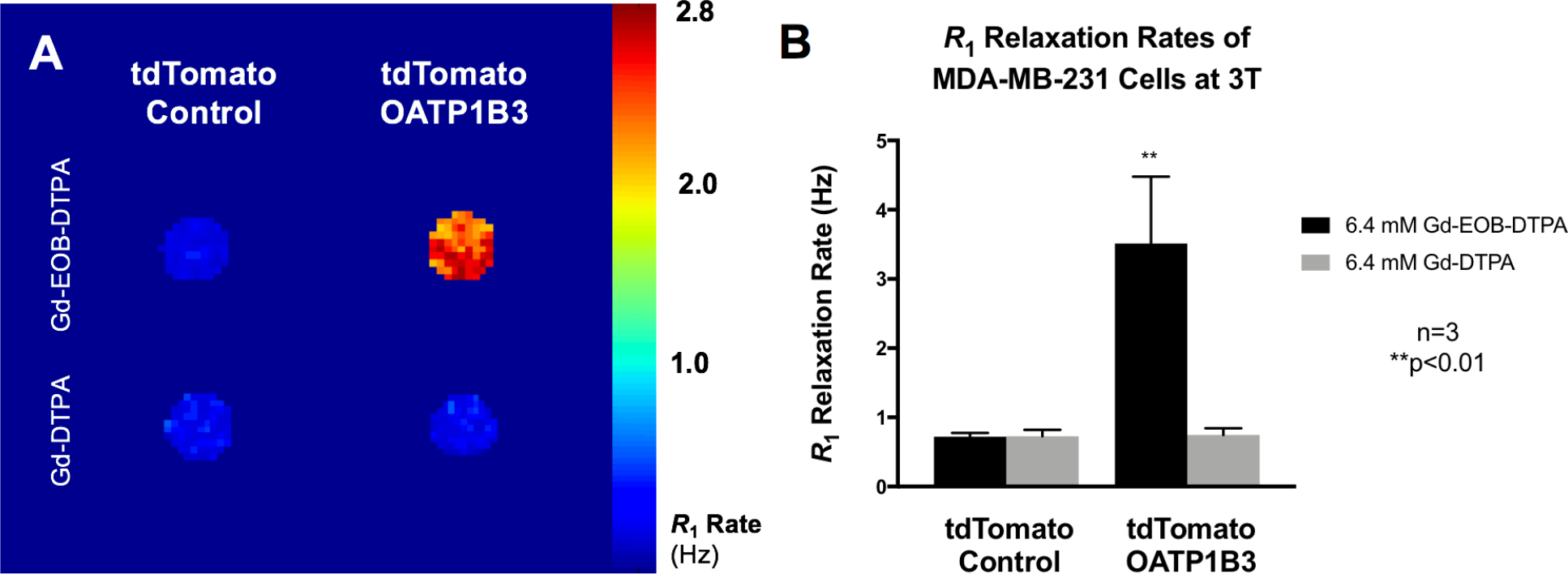
Spin-Lattice Relaxation Rates of Engineered MDA-MB-231 Cells at 3 Tesla. A) Spin-lattice relaxation map of tdTomato Control and tdTomato OATP1B3 cells treated with 6.4-mM Gd-EOB-DTPA or 6.4-mM Gd-DTPA. B) Spin-lattice relaxation rates (*R*1, Hz) of tdTomato Control and tdTomato OATP1B3 cells treated with 6.4-mM Gd-EOB-DTPA or 6.4-mM Gd-DTPA. Error bars represent one standard deviation.

**Supplementary Figure 3.**
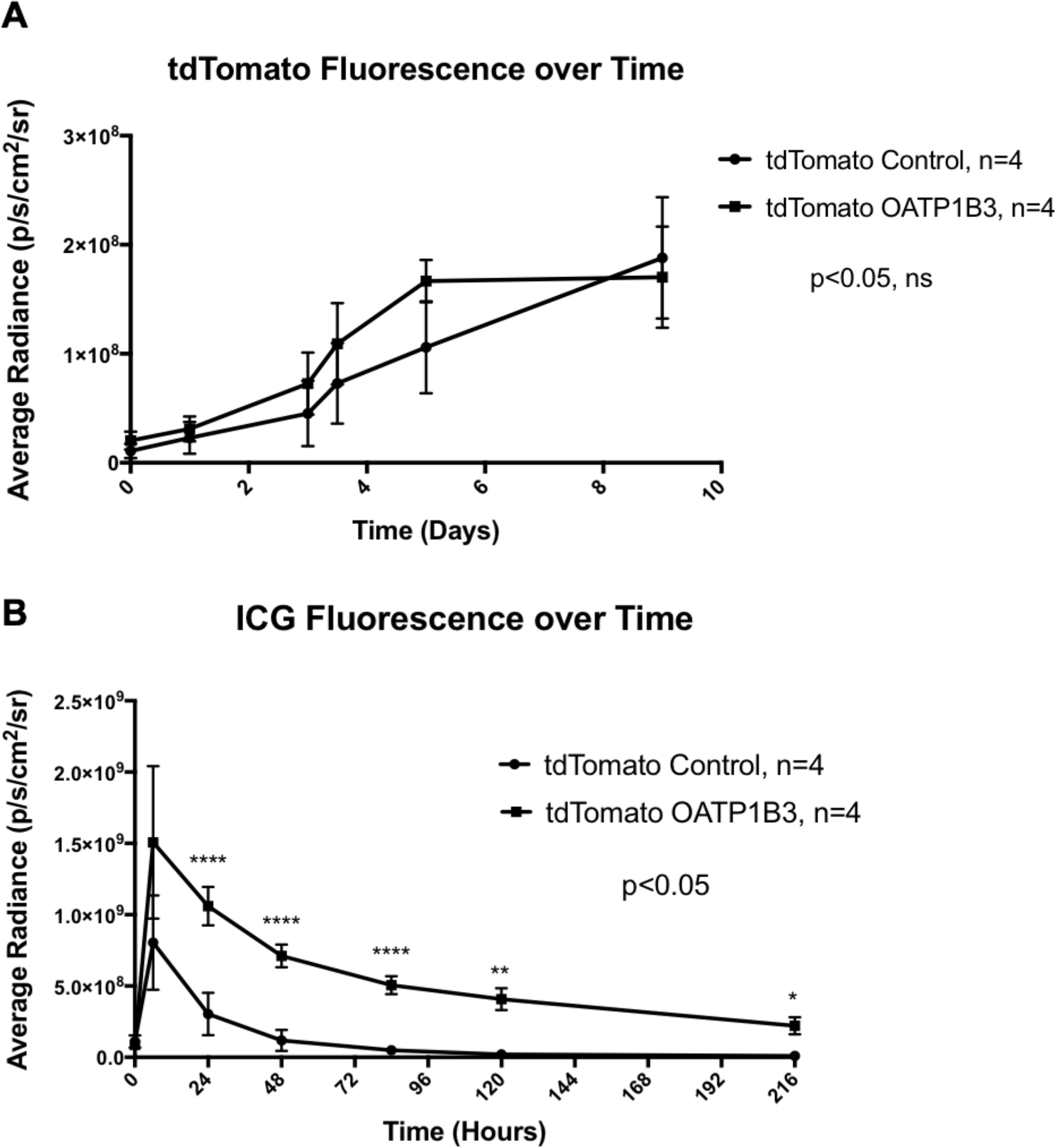
Longitudinal *In Vivo* Fluorescence Imaging of Engineered Tumors. A) tdTomato fluorescence radiance (p/s/cm^2^/sr) of tdTomato Control and tdTomato OATP1B3 tumors over time, up to 9 days. B) ICG fluorescence radiance (p/s/cm^2^/sr) of tdTomato Control and tdTomato OATP1B3 tumors over time, up to 9 days, following administration of 8 mg/kg ICG. Error bars represent one standard deviation.

## MATERIALS AND METHODS

### Lentiviral Construction and Production

Genetic engineering of the reporter gene system involved third-generation packaging and envelope-expression plasmids (pMDLg/pRRE, pRSV-Rev, and pMD2.G, Addgene plasmids: #12251, #12253, and #12259, respectively; gifts from Didier Trono). A lentiviral transfer vector encoding the *tdTomato* (*tdT*) reporter gene for fluorescence under the human elongation factor 1 alpha promoter (pEF1α) was acquired, along with a second lentiviral transfer plasmid encoding the *zsGreen1* (*zsG*) reporter gene for fluorescence and the *Organic anion transporting polypeptide 1a1* (*Oatp1a1*) reporter gene separated via a P2A self-cleaving peptide, also under regulation of pEF1α^61^. The sequence for *Organic anion transporting polypeptide 1b3* (*Oatp1b3*) was acquired from the hOATP1B3/SLCO1B3 VersaClone cDNA Vector (Cat. RDC0870, R&D Systems, Minneapolis, Minnesota, United States). Cloning was performed using In-Fusion HD Cloning (Takara Bio USA Inc, Madison, Wisconsin, United States). The *Oatp1a1* sequence in the second plasmid was replaced by the *Oatp1b3* sequence to obtain a resultant zsG/Oatp1b3 lentiviral transfer plasmid. The *zsG* fluorescent protein was selected for fluorescence-activated cell sorting of *Oatp1b3*-expressing cells, as it absorbs virtually no near-infrared light within the 680 to 970 nm wavelength range and thus, will not confound signals generated during analysis of *Oatp1b3* as a reporter gene for NIR-PAI^62^. To produce tdT and zsG/Oatp1b3 lentiviruses (LV-tdT and LV-zsG/Oatp1b3), the packaging, envelope and one of these transfer plasmids were co-transfected into human embryonic kidney (HEK 293T) cells using Lipofectamine 3000 according to the manufacturer’s lentiviral production protocol (Thermo Fisher Scientific Inc., Waltham, Massachusetts, United States). Lentivirus-containing supernatants were harvested 24h and 48h post transfection, filtered through a 0.45-μm filter, and stored at −80°C prior to use.

### Cell Culturing and Generation of Stable Cells

Human embryonic kidney cells (HEK 293T), human triple negative breast cancer cells (MDA-MB-231), and murine triple negative breast cancer cells (4T1) were obtained from a commercial supplier (American Type Culture Collection; ATCC, Manassas, Virginia, United States) and cultured in Dulbecco’s Modified Eagle Media (DMEM, Wisent Inc., Saint-Jean-Baptiste, Quebec, Canada) supplemented with 10% fetal bovine serum at 37°C and 5% CO_2_. Cells were routinely tested negative for mycoplasma using the MycoAlert mycoplasma detection kit (Lonza Group, Basel, Switzerland). MDA-MB-231 and 4T1 cells were transduced with LV-tdT overnight in the presence of 4 to 8-μg/mL polybrene. Transduced cells were washed, collected, and sorted using a FACSAria III fluorescence-activated cell sorter (BD Biosciences, Mississauga, Ontario, Canada) for positive tdTomato fluorescence, generating tdTomato Control cells. A subset of these tdTomato Control cells was then transduced a second time with LV-zsG/Oatp1b3 and sorted for equivalent tdTomato fluorescence intensity relative to the original tdTomato Control cells, as well as for positive zsGreen fluorescence. The resulting population was named tdTomato OATP1B3 cells. Sorted cells were utilized for all subsequent *in vitro* and *in vivo* experiments.

### Immunofluorescence Staining

Cells were grown on glass coverslips, rinsed with PBS, and fixed with 4% paraformaldehyde (PFA) for 10 minutes. Following this, cells were permeabilized via 0.02% Tween 20 for 20 minutes and incubated overnight at 4°C with rabbit anti-SLCO1B3 (OATP1B3) primary antibody (2-μg/mL working concentration, HPA004943, Sigma-Aldrich Canada, Oakville, Ontario, Canada). Goat anti-rabbit AlexaFluor 647-conjugated secondary antibody was then applied for two hours at room temperature (1:500 dilution; 4-μg/ml working concentration, ab150079, Lot E114795, Abcam, Cambridge, Massachusetts, United States). Coverslips were counterstained with DAPI and imaged using an LSM Meta 510 microscope (Carl Zeiss AG, Oberkochen, Germany).

### Proliferation Assays

Non-transduced, tdTomato Control, and tdTomato OATP1B3 cells (5×10^4^) were seeded in 96-well plates and cell proliferation was evaluated using a tetrazolium salt (3-(4,5-dimethylthiazol-2-yl)-2,5-diphenyltetrazolium bromide, MTT) assay. Cells were incubated in phenol red-free DMEM (Cat. 319-051-CL, Wisent Inc., Saint-Jean-Baptiste, Quebec, Canada) supplemented with 10% FBS that contained either 35-μg/ml ICG or an equivalent volume of dimethyl sulfoxide (DMSO), which is used as the solvent for the stock ICG solution. Prior to optical measurements, wells were washed three times with PBS containing DMSO (10% v/v). Absorbance measurements at 590 nm were acquired with a spectrophotometer at 0, 24 and 48 hours after seeding. Proliferative measurements at 24 and 48 hours were normalized to absorption values obtained at seeding (0 hours).

### *In Vitro* Fluorescence and Photoacoustic Imaging

Human (MDA-MB-231) and murine (4T1) tdTomato Control and tdTomato OATP1B3 cells (2×10^6^) were seeded and grown for three days in 150mm-diameter dishes (Cat. SIAL0599, Sigma-Aldrich Canada, Oakville, Ontario, Canada). Cells were incubated for 90 minutes at 37°C and 5% CO_2_ in media containing either 35-μg/ml ICG or an equivalent volume of DMSO solvent. Cells were then washed three times with DMSO-containing PBS (10% v/v), trypsinized, counted and 3×10^6^ cells were pelleted and embedded into a 25-μ;L volume of 1% agarose, 0.5% intralipid solution. The pellets were placed on ice for 5 minutes, allowing the gel to solidify, thus generating a small, agarose-embedded cell sphere for each engineered cell and treatment group. Each cell sphere was subsequently transferred into the wells of a custom-built 1% agarose, 0.5% intralipid phantom, designed to be asymmetrical in its distribution of wells. Additionally, a sphere containing 35-μg/ml ICG, and a gel sphere containing media without ICG were transferred to the phantom as positive and negative controls, respectively. FLI of the phantom was performed via an IVIS Lumina XRMS *In Vivo* Imaging System (PerkinElmer, Waltham, Massachusetts, United States) to measure ICG fluorescence intensity using a 0.5-s exposure time, 780-nm excitation filter, and 845-nm emission filter. A region of interest (ROI) was drawn around the perimeter of each well. Radiance values for ICG (p/s/cm^2^/sr) were measured from each ROI using LivingImage software (PerkinElmer, Waltham, Massachusetts, United States). Following FLI, the phantom wells were immediately sealed with 1% agarose, 0.5% intralipid solution and allowed to gel prior to PAI.

Following FLI, wells in the phantom were backfilled with a 1% agarose, 0.5% intralipid solution. PAI was then performed using a custom photoacoustic tomography system. The system consisted of a transducer array with 28 custom-built cylindrical unfocused transducers (2.7-MHz central frequency, ~127% bandwidth, 4.5-mm diameter) mounted on two concentric circular rungs. All transducers were angled inwards to a point 25 mm below the array, providing a field-of-view ~20 mm in diameter. 780-nm NIR illumination was provided by a tunable (680 – 950 nm) 10-Hz-pulsed laser (Phocus InLine, Opotek Inc., California, United States) injected into a four-legged fused-end fiber bundle (Lumen Dynamics Group Inc., Mississauga, Ontario, Canada). The output ends of the fiber bundle were positioned at the centre of the circular transducer rings, pointing inwards towards the same focal point as the transducers. The transducer array with attached fiber bundles was mounted to an Epson SCARA robot (Epson C4, Suwa, Nagano Prefecture, Japan) for scanning. All transducers were connected to a custom 28-channel, 50-MHz data acquisition system. The cell phantom was sealed with water in a modified Ziploc sandwich bag (S. C. Johnson & Son, Racine, Wisconsin, United States) and submerged in a water tank along with the transducer array. The imaging array was raster scanned in three dimensions with 4-mm steps in the XY-plane, and 5-mm steps in the Z-direction to image the entire phantom. At each scan point, signal averaging was performed over 16 laser pulses to increase signal-to-noise ratio. Following imaging, image reconstruction was performed using Universal Back-projection^63^. A maximum intensity projection (MIP) of the 3D stack was then acquired using ImageJ^64^. For each phantom (n=3), photoacoustic MIPs were scaled and overlayed on annotated fluorescence images and ROIs from FLI were transferred to photoacoustic MIPs. Average signal intensity was measured from each ROI and contrast-to-noise ratio (CNR) was calculated relative to an equal-sized ROI positioned at the centre of the MIP.

### *In Vitro* Magnetic Resonance Imaging at 3 Tesla

Non-transduced, tdTomato Control, and tdTomato OATP1B3 cells (2×10^6^) were seeded in T-175 flasks and grown for 3 days. Cells were incubated with media containing 6.4-mM Gd-EOB-DTPA (Eovist/Primovist®, Bayer Health Care Pharma, Berlin, Germany) or Gd-DTPA (Magnevist®, Bayer Schering Pharma, Berlin, Germany) for 90 minutes at 37°C and 5% CO_2_. Cells were then washed three times with PBS, trypsinized and pelleted in 0.2-mL tubes, and placed into a 2% agarose phantom mold that was incubated in a 37°C chamber for two hours to mimic body temperature. MRI was performed on a 3-Tesla GE clinical MR scanner (General Electric Healthcare Discovery MR750 3.0 T, Milwaukee Wisconsin, United States) and a 3.5-cm diameter birdcage RF coil (Morris Instruments, Ottawa, Ontario, Canada). A fast spin echo inversion recovery pulse sequence was used with the following parameters: field of view = 256×256, repetition time (TR) = 5000 ms, echo time = 19.1 ms, echo train length = 4, number of excitations = 1, receiver bandwidth = 12.50 MHz, inversion times (TI) = 20, 35, 50, 100, 125, 150, 175, 200, 250, 350, 500, 750, 1000, 1500, 2000, 2500, 3000, in-plane resolution = 0.27 mm^2^, slice thickness = 2.0 mm. Spin-lattice relaxation rates were computed via MatLab (R2018a, MathWorks, Natick, Massachusetts, United States) by overlaying the image series and calculating the signal intensity on a pixel-by-pixel basis across the inversion time image series, followed by a fitting of the data into the following equation to output the spin-lattice relaxation time (*T*1), where 𝑆 represents signal intensity, *κ* represents the scaling factor, and *ρ* represents proton spin density:

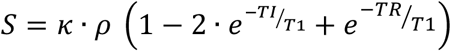

### Orthotopic Tumor Model

Animal experiments were undertaken with an approved protocol of the University of Western Ontario’s Council on Animal Care (AUP 2016-026) and in accordance with the standards of the Canadian Council on Animal Care. One million tdTomato Control or tdTomato OATP1B3 MDA-MB-231 cells were implanted orthotopically into the right-bearing fourth mammary fat pad of immunocompromised (NU-*Foxn1*^nu^) 6-8-week-old female mice (Charles River Laboratories, Wilmington, Massachusetts, United States). Forty-eight hours post-cell implantation, fluorescence and photoacoustic images of mice were collected pre-contrast and 24 hours post-intraperitoneal administration of 8-mg/kg ICG. As described below, fluorescence images using a 520-nm excitation and 570-nm emission filter were collected to evaluate *tdTomato* expression as a surrogate measure of cell number for both tdTomato Control and tdTomato OATP1B3 tumors in order to normalize fluorescence data for ICG with tumor size.

### *In Vivo* Fluorescence Imaging

FLI was performed on an IVIS Lumina XRMS *In Vivo* Imaging System (PerkinElmer, Waltham, Massachusetts, United States). Mice were anesthetized with 1-2% isofluorane using a nose cone attached to an activated carbon charcoal filter for passive scavenging. FLI for tdTomato was performed using 520-nm excitation and 570-nm emission filters and images were acquired with a 5-minute exposure time. FLI for ICG was performed using 780-nm excitation and 845-nm emission filters and images were acquired with a 5-min exposure time. ROIs were manually drawn around tumor borders on tdTomato images using LivingImage software (PerkinElmer, Waltham, Massachusetts, United States) to measure tdTomato fluorescence radiance (p/s/cm^2^/sr), and then the ROI for each mouse was copied to the ICG image to measure ICG fluorescence radiance (p/s/cm^2^/sr). The ratio of ICG to tdTomato fluorescence, referred to as “normalized fluorescence” in accordance with previous literature^65^, was then calculated for each tumor both prior-to and 24 hours following ICG administration.

### *In Vivo* Photoacoustic Imaging

Mice were anesthetized as for FLI, and US and PAI was performed using the Vevo LAZR-X System (Fujifilm VisualSonics Inc., Toronto, Ontario, Canada). An LZ550 transducer (Fujifilm VisualSonics Inc., Toronto, Ontario, Canada) was used to acquire US and PAI images with axial resolutions of 40 μm. US images of tumors were first acquired, followed by NIR spectrum (680-970 nm) PAI in 5-nm increments. For each tumor, a central slice, along with adjacent slices 800 μ;m on either side were acquired. The liver of one mouse was imaged (n=1) immediately following intravenous administration of ICG (8 mg/kg) to generate a positive control PAI spectrum for intracellular *in vivo* ICG. Vevo Lab Image Software (Fujifilm VisualSonics Inc., Toronto, Ontario, Canada) was used to outline ROIs around the perimeter of each tumor in ultrasound images, and these ROIs were subsequently overlayed onto corresponding photoacoustic images. Tumor ROIs were processed through the software to generate a full-spectrum photoacoustic plot for each tumor both before and 24 hours following administration of ICG. ICG signals were spectrally unmixed from tumor data to generate ICG localization images within tumors, as previously described^33^.

### *Ex Vivo* FLI and Histology

Following *in vivo* imaging, mice were immediately sacrificed via isofluorane overdose, perfused with 4% paraformaldehyde through the left heart ventricle and tumors were excised from mammary fat pads. Tumors were subsequently frozen in Tissue-Tik Optimum Cutting Temperature (OCT) medium (Sakura Finetek, Maumee, Ohio, United States) and both 150-μ;m and 10-μ;m frozen sections were collected via the Leica CM350 Cryostat (Leica Microsystems, Wetzlar, Germany). The 150-μ;m sections were plated on glass slides and FLI images were collected using the IVIS Lumina XRMS *In Vivo* Imaging System (PerkinElmer, Waltham, Massachusetts, United States) for tdTomato expression (5-s exposure, 520-nm excitation filter, 570-nm emission filter), zsGreen expression (10-s exposure, 480-nm excitation filter, 520-nm emission filter), and ICG uptake/retention (10-s exposure, 780-nm excitation filter, 845-nm emission filter). Microscopy images for tdTomato and zsGreen fluorescence were taken of the 10-μ;m sections using an EVOS FL Auto 2 Imaging System (Invitrogen, Thermo Fisher Scientific Inc., Waltham, Massachusetts, United States).

### Statistics

One-way Analysis of Variance (ANOVA) was performed followed by Tukey’s post-hoc multiple comparisons using Graphpad Prism software (Version 7.00 for Mac OS X, GraphPad Software Inc., La Jolla, California, United States, www.graphpad.com) for cell proliferation studies, *in vitro* FLI, PAI and MRI, and *in vivo* FLI and PAI. For all tests, a nominal p-value less than 0.05 was considered statistically significant.

